# High-Dimensional Immunophenotyping Identifies Circulating Immune Signatures Associated with FeNO-Defined Asthma Phenotypes

**DOI:** 10.64898/2026.09.04.749390

**Authors:** Neha Solanki, Nick Wanner, James A. Thomas, William Toth, Sobia Farooq, Yifan Wang, Serpil Erzurum, Kewal Asosingh

**Author notes:** **Corresponding Author:** Neha Solanki, MD, 2049 E 100th Street A Building.

## Abstract

**Background:** Asthma is a heterogeneous inflammatory airway disease traditionally classified into T2-high (eosinophilic) and T2-low inflammatory phenotypes. Fractional exhaled nitric oxide (FeNO) is commonly used as a biomarker of T2 inflammation, although nitric oxide also regulates monocyte maturation and T-cell polarization. We hypothesized that FeNO identifies distinct systemic immune phenotypes in asthma.

**Methods:** Peripheral immune-cell subsets were characterized using OMIP-69, a 40-color high-dimensional immunophenotyping panel encompassing innate and adaptive immune populations. Adults with asthma and healthy controls underwent peripheral blood immunophenotyping. FeNO was dichotomized at 25 ppb (FeNO-high ≥25 vs FeNO-low <25). Immune-cell subset proportions were summarized as medians (IQRs) and compared using the Wilcoxon rank-sum test. Logistic regression was used to evaluate associations between immune subsets and FeNO status.

**Results:** Eighteen participants with asthma and seven healthy controls were enrolled. Asthmatic participants had higher BMI than healthy controls (median [IQR] asthma: 35.1 [28.9– 37.7] vs healthy: 25.4 [22.6–26.7] kg/m²; p=0.003), although demographics were otherwise similar. Among asthmatics, 10 participants had FeNO-low asthma, and eight had FeNO-high asthma. FeNO-high asthma demonstrated increased classical monocytes (CD14++CD16−) and compared with healthy controls (p=0.027), whereas FeNO-low asthma more closely resembled healthy controls. Intermediate monocytes (CD14++CD16+) were increased in FeNO-high versus FeNO-low (p= 0.08); a similar association was noted for classical monocytes (p = 0.13). In contrast, CCR7+ γδ T cells were significantly enriched in FeNO-low asthma (p=0.002). Logistic regression demonstrated positive associations between high FeNO and classical monocytes (OR 16.06, 95% CI 1.56–1304.69; p=0.007) and intermediate monocytes (OR 4.19, 1.22–132.30; p=0.017), whereas CCR7+ γδ T cells were negatively associated with high FeNO (OR 0.06, 0.00–0.47; p=0.001).

**Conclusions:** Distinct systemic immune signatures identified by high-dimensional cytometry support the existence of biologically divergent FeNO-defined asthma endotypes and may inform future biomarker-driven approaches to patient stratification and precision therapy.

**Graphical Abstract:** 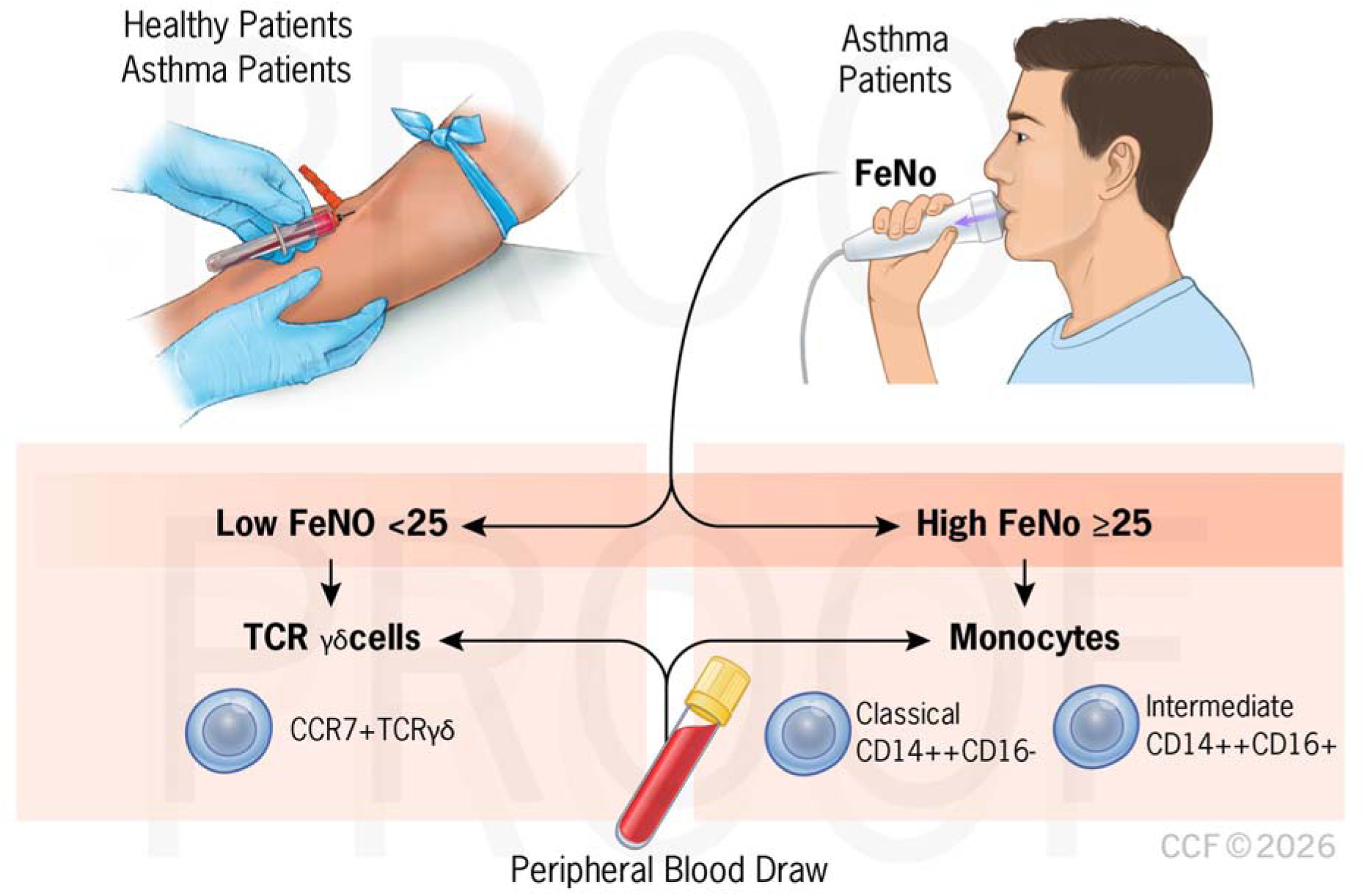

## Introduction

Asthma is a fundamentally heterogeneous disease with diverse underlying mechanisms that have significant implications for diagnosis, treatment, and outcomes. Asthma phenotypes represent clusters of demographic, clinical, and pathophysiological characteristics in patients. Endotypes are distinct disease subsets defined by specific biologic pathways, including differences in inflammatory mechanisms, genetic predisposition, and immune responses. The most important endotypes are T2-high and T2-low asthma. T2-high asthma is characterized by an enhanced Th2 immune response: high serum levels of IL-4, IL-5, and IL-13, and elevated total IgE. Clinical biomarkers are also elevated: blood eosinophils > 300 µL and fractional exhaled nitric oxide (FeNO) ≥ 25 parts per billion (ppb). T2-low asthma comprises subtypes that are under characterized, including T17-high asthma, T2-low, and T2-low/T17-low asthma. Understanding of T2-low asthma is limited because it is defined by the absence of T2 or Type 2-high inflammatory markers. In severe asthma, we have phenotype-guided treatment, specifically for T2-high asthma. There is a lack of effective therapies for T2-low and T2-mixed asthma, reflecting an incomplete understanding of the underlying mechanisms ^1,2^.

Nitric oxide (NO) is a gas produced in the airways by inducible nitric oxide synthase 2 (iNOS2) enzyme, which is upregulated in the airway epithelium during T2 inflammation^3^. FeNO correlates with airway eosinophilic inflammation, specifically reflecting IL-13 driven upregulation of inducible nitric oxide synthase, and provides a noninvasive surrogate for assessing T2 airway inflammation ^3,4^. FeNO is not diagnostic of asthma but is used as a tool alongside a patient’s history, physical examination, spirometry, and biomarker data ^5^. Elevated FeNO levels are associated with a patient’s increased responsiveness to therapy targeting IL-4 and IL-13 pathways, highlighting the relationship between airway nitric oxide production and type 2 immune activity. Therefore, FeNO reflects a broader immunologic process that extends beyond airway epithelial biology.

By using known biomarkers for T2-high asthma, clinicians aim to diagnose, manage, and treat the condition. Many individuals remain uncontrolled despite adherence to inhaler therapy and advanced biologic treatment because of our limited knowledge and binary view of asthma endotyping. While flow cytometry–based endotyping has transformed disease classification in oncology and autoimmunity, asthma phenotyping remains largely biomarker-driven (e.g., FeNO, eosinophils), with limited integration of high-dimensional immune cell states. Deep immunophenotyping yields a precise endotype understanding and reveals how circulating cell populations are amenable to targeted therapy, particularly for T2-low asthma ^2,4,6,7^. Understanding whether specific circulating cell populations differ by FeNO status could illuminate the immunologic mechanisms driving different asthma endotypes, linking airway inflammation with systemic immune regulation ^8,9^.

In this study, we utilized high-dimensional flow cytometry to define the systemic immune landscape of asthma. We hypothesized that FeNO-high and FeNO-low asthma represent biologically distinct inflammatory endotypes characterized by unique circulating immune cell signatures. To our knowledge, this high-dimensional immunophenotyping panel has not previously been used to investigate asthma. By stratifying individuals according to FeNO levels, we sought to identify immune profiles associated with FeNO-defined phenotypes and gain insight into the mechanisms underlying asthma heterogeneity. The data showed that high-dimensional immunophenotyping identified distinct systemic immune signatures associated with FeNO-defined asthma phenotypes. FeNO-high asthma was characterized by monocyte enrichment, whereas FeNO-low asthma demonstrated increased CCR7+ γδ T cells.

## Materials and Methods

### Sample Collection

Adult (≥ 18 years old) healthy controls and asthma patients from the Cleveland Clinic were recruited to this study from 2023-2025. Asthmatics had received the diagnosis from the Cleveland Clinic Asthma Center based on their symptoms and medical history. Healthy controls and asthmatics had no diagnosis of cancer or autoimmune disorder. Smokers were excluded from the study. Healthy controls had no eczema, allergies, or other Type 2-mediated disease. Asthmatics had no other pulmonary diagnosis such as COPD or bronchiectasis. The study was conducted under approval from the **Cleveland Clinic Institutional Review Board (IRB Protocol #26-077)**, and all participants had previously provided written informed consent for study participation.

### Flow Cytometry

Blood was collected in EDTA tubes and processed using Ficoll density centrifugation to isolate mononuclear immune cells for downstream analysis. Peripheral blood mononuclear cells (PBMCs) were cryopreserved immediately after isolation and then thawed in batches for subsequent staining with fluorophore-conjugated antibodies, fixation, and permeabilization along with an aliquot of control cells to control for batch effects.

A 40-color spectral flow cytometry panel based on OMIP-069 was used (Supplementary Table I). The staining protocol described in the OMIP-069 methods publication was followed and applied to asthmatics and healthy individuals ^10,11^. Following staining, cells were washed and resuspended for acquisition on a 5-laser Cytek Aurora spectral cytometer (Cytek Biosciences, Fremont, CA). Spectral unmixing was performed with single-stained reference controls using PBMCs. A minimum of 100,000 events were acquired per sample to ensure adequate detection of rare immune cell populations.

Flow cytometry data were analyzed using FlowJo Version 10 (FlowJo, LLC, Ashland, OR) and FCS Express Version 7 (De Novo Software, Pasadena, CA). Events were gated against time to exclude periods of fluidic instability, followed by sequential gating to exclude debris, doublets, and dead cells. Subsequent analyses were performed on CD45+ leukocytes using standard gating strategies to identify major immune cell subsets including CD4 T cells, CD8 T cells, regulatory T cells, γδ T cells, NKT-like cells, B cells, NK cells, monocytes, and dendritic cells ^10,11^.

Antibodies were purchased from BD Biosciences (Franklin Lakes, NJ), BioLegend (San Diego, CA), Miltenyi Biotec (Bergisch Gladbach, Germany), Thermo Fisher Scientific (Waltham, MA), and Cytek Biosciences (Fremont, CA) as indicated in the reagent table. Reagents were purchased as reflected in the OMIP-069 panel (Supplemental Table 1).

### Data Analysis

Demographic and clinical characteristics were compared between healthy controls and participants with asthma using the Wilcoxon rank-sum test for continuous variables and chi-square or Fisher’s exact test, as appropriate, for categorical variables. Continuous variables are presented as median [interquartile range (IQR)], and categorical variables as counts and percentages.

Fractional exhaled nitric oxide (FeNO) was measured according to standardized American Thoracic Society guidelines, with measurements obtained prior to spirometry while participants exhaled at a constant flow rate. FeNO was dichotomized using the clinically relevant threshold of 25 ppb (FeNO-low <25 ppb vs. FeNO-high ≥25 ppb), consistent with American Thoracic Society recommendations identifying patients in whom type 2 airway inflammation and corticosteroid responsiveness are more likely.

High-dimensional cytometry is exploratory and hypothesis-generating^12^. Screening immune phenotypes and carrying candidate populations forward for clinical correlations is an accepted discovery workflow^13^. Because this was an exploratory immunophenotyping study with a limited sample size, immune cell subsets were first screened for differences between participants with asthma and healthy controls using Wilcoxon rank-sum tests. Broad parent populations (lymphocytes and monocytes) were included to provide gating context but were not emphasized in the interpretation because these populations were subsequently resolved into more specific immune subsets.

Given the exploratory nature of high-dimensional immunophenotyping studies, immune subset analyses were performed to identify candidate FeNO-associated immune signatures^12,14^. Similar discover-based immune-profiling strategies have been used to characterize circulating immune phenotypes associated with clinical outcomes^15^. Overall group differences were assessed using Kruskal-Wallis testing, with Benjamini-Hochberg FDR adjustment performed across evaluated immune subsets^12^. Candidate immune populations identified from this exploratory screen were subsequently evaluated using Spearman rank correlation coefficients to determine whether subset frequencies demonstrated biologically consistent relationships with continuous FeNO levels^16^. Similar approaches have utilized Spearman correlation analyses to characterize relationships between immune cell populations and clinically relevant phenotypes. Candidate FeNO-associated immune subsets were further characterized using Dunn’s post-hoc comparisons to evaluate differences between healthy controls, FeNO-low asthma, and FeNO-high asthma groups. Given the exploratory nature of the study, immune subsets demonstrating nominal associations were prioritized for subsequent exploratory biological characterization across FeNO-defined phenotypes^17^.

Candidate FeNO-associated immune cell subsets identified through exploratory immunophenotyping were further evaluated using Firth penalized logistic regression to estimate associations with FeNO-high asthma status. Firth regression was selected because of the limited sample size and potential for separation in logistic regression models^18–20^. Odds ratios with 95% confidence intervals were calculated to estimate the association between immune subset frequencies and FeNO-high asthma status. For Firth logistic regression, immune subset frequencies were standardized prior to modeling, and odds ratios are reported per 1 standard deviation increase. Confidence intervals were generated using profile penalized likelihood methods. To evaluate whether findings were influenced by individual observations, leave-one-out sensitivity analyses were performed by sequentially excluding each participant and refitting the model. Results were summarized using forest plots.

All tests were two-sided. Both nominal and Benjamini-Hochberg-adjusted p-values are reported, with statistical significance in focused analyses defined as adjusted p < 0.05. Missing data were handled using complete case analysis for each statistical comparison. No imputation was performed.

## Results

### Clinical Characteristics

We recruited 7 healthy controls and 18 individuals with asthma. There were no significant demographic differences between the groups. However, individuals with asthma had a higher body mass index (BMI) compared with healthy controls (median [IQR]: 35.1 kg/m² [28.9–37.8] vs 25.4 kg/m² [22.6–26.7], p = 0.003) (Table 1). Sex, age, and race were similar between groups (p > 0.05) (Table 2).

**Table 1.**
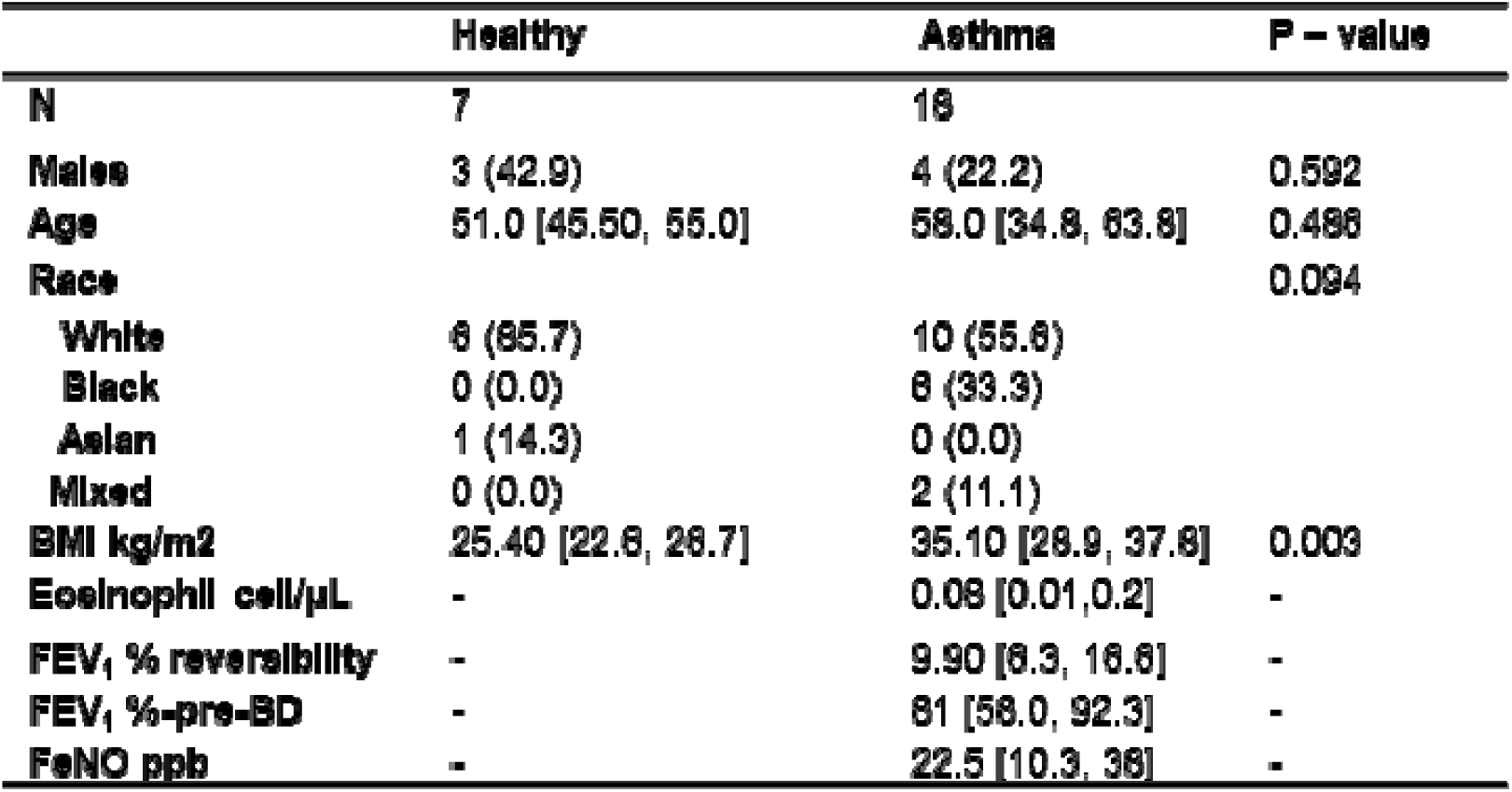
Baseline demographic and clinical characteristics of study participants. Participant characteristics are summarized for healthy controls (n = 7) and individuals with asthma (n = 18). Continuous variables are presented as median [IQR] and categorical variables as n (%). P-values compare groups using appropriate statistical tests. Abbreviations: BMI = body mass index; μL= microliter; FEV1 = Forced expiratory volume in 1 second; FeNO: fractional excretion of nitric oxide; ppb = parts per billion.

**Table 2.**
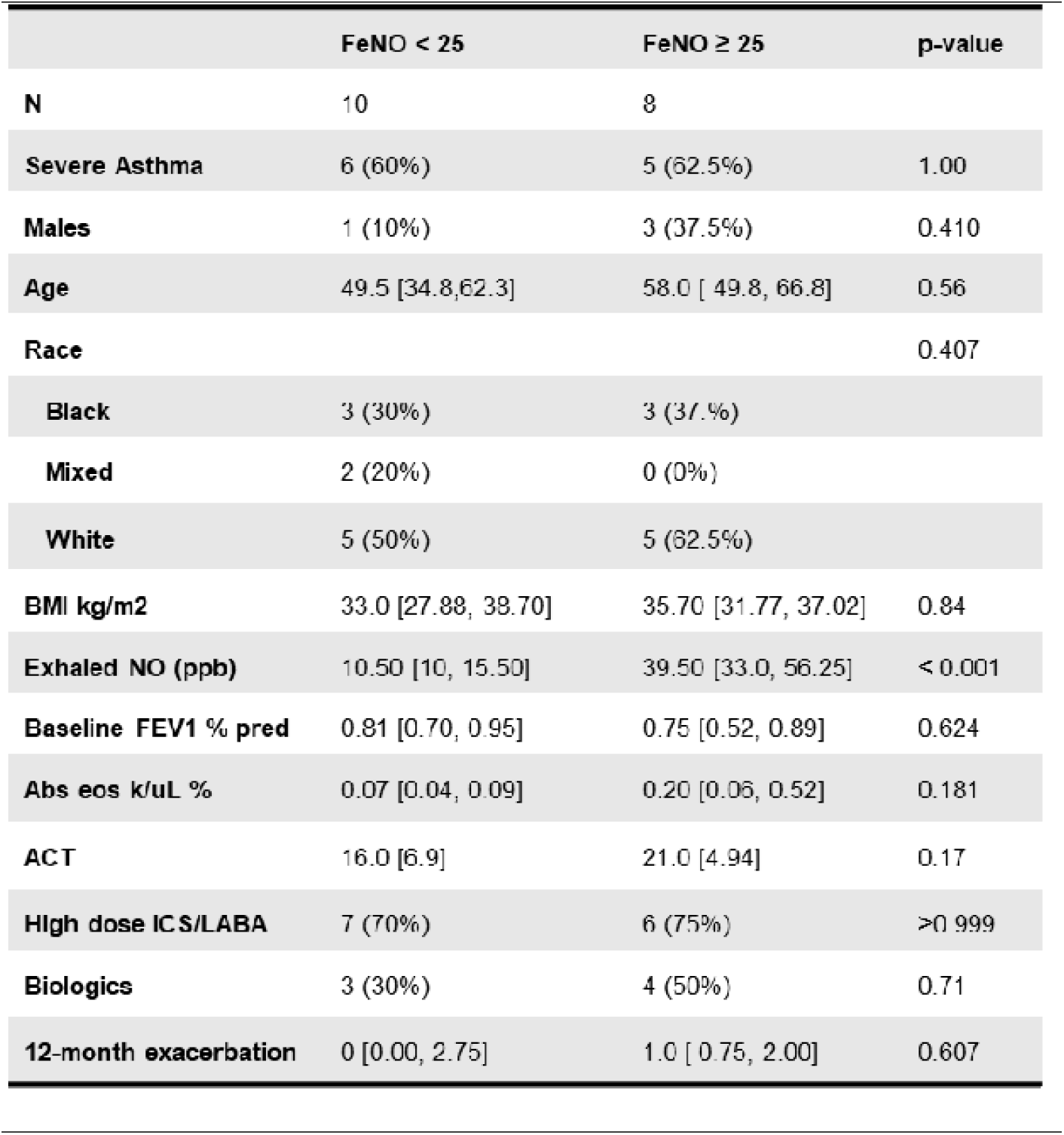
Baseline demographic and clinical characteristics stratified by FeNO category (<25 vs ≥25 ppb). Continuous variables are presented as median (IQR) or mean (SD) as appropriate; categorical variables are presented as n (%). Group comparisons were performed using Wilcoxon rank-sum tests or χ²/Fisher’s exact tests. Baseline demographic and clinical characteristics were similar between participants with low (<25 ppb) and high (≥25 ppb) FeNO. Abbreviations: ACT = asthma control test; ICS/LABA = inhaled corticosteroids and long-acting beta agonist.

|  | FeNO < 25 | FeNO ≥ 25 | p-value |
| --- | --- | --- | --- |
| **N** | 10 | 8 |  |
| **Severe Asthma** | 6 (60%) | 5 (62.5%) | 1.00 |
| **Males** | 1 (10%) | 3 (37.5%) | 0.410 |
| **Age** | 49.5 [34.8, 62.3] | 58.0 [ 49.8, 66.8] | 0.56 |
| **Race** |  |  | 0.407 |
| **Black** | 3 (30%) | 3 (37.5%) |  |
| **Mixed** | 2 (20%) | 0 (0%) |  |
| **White** | 5 (50%) | 5 (62.5%) |  |
| **BMI kg/m2** | 33.0 [27.88, 38.70] | 35.70 [31.77, 37.02] | 0.84 |
| **Exhaled NO (ppb)** | 10.50 [10, 15.50] | 39.50 [33.0, 56.25] | < 0.001 |
| **Baseline FEV1 % pred** | 0.81 [0.70, 0.95] | 0.75 [0.52, 0.89] | 0.624 |
| **Abs eos k/uL %** | 0.07 [0.04, 0.09] | 0.20 [0.06, 0.52] | 0.181 |
| **ACT** | 16.0 [6.9] | 21.0 [4.94] | 0.17 |
| **High dose ICS/LABA** | 7 (70%) | 6 (75%) | >0.999 |
| **Biologics** | 3 (30%) | 4 (50%) | 0.71 |
| **12-month exacerbation** | 0 [0.00, 2.75] | 1.0 [ 0.75, 2.00] | 0.607 |

Participants with asthma demonstrated a non-eosinophilic profile, with low absolute eosinophil counts (AEC: 0.08 cells/µL [0.00–0.16]) and relatively low FeNO levels (FeNO: 22.5 ppb [10.3–38.0]) (Table 1). Lung function was generally preserved, with a median percent predicted forced expiratory volume in 1 second (FEV₁% predicted) of 81.0% [58.0–92.3] (Table 1). 72.2% of asthmatics were on an inhaled corticosteroid and long-acting beta-agonist and 38.9% of asthmatics were on a biologic agent at that time of study recruitment. Given the heterogeneity of airway inflammation in asthma, we next stratified individuals with asthma based on FeNO levels to examine whether immune signatures differed according to airway nitric oxide levels. Participants were categorized into FeNO-low (n = 10) and FeNO-high (n = 8) groups using a threshold of 25 ppb. As expected, FeNO levels differed significantly between the groups (FeNO-low: 10.5 ppb [10.0–15.5] vs FeNO-high: 39.5 ppb [33.0–56.3], p < 0.001). However, there were no clinical differences between the two groups and there was no difference in medications between the two groups.

### Circulating Immunophenotypes in FeNO-High and FeNO-Low Asthmatics and Healthy Controls

This project utilized a hierarchical gating strategy for an established and previously validated 40-color multiparametric panel^10,11^ (Figure 1). This 40-color panel characterizes CD4 T cells, CD8 T cells, regulatory T cells, γδ T cells, NKT-like cells, B cells, NK cells, monocytes, and dendritic cells.

**Figure 1.**
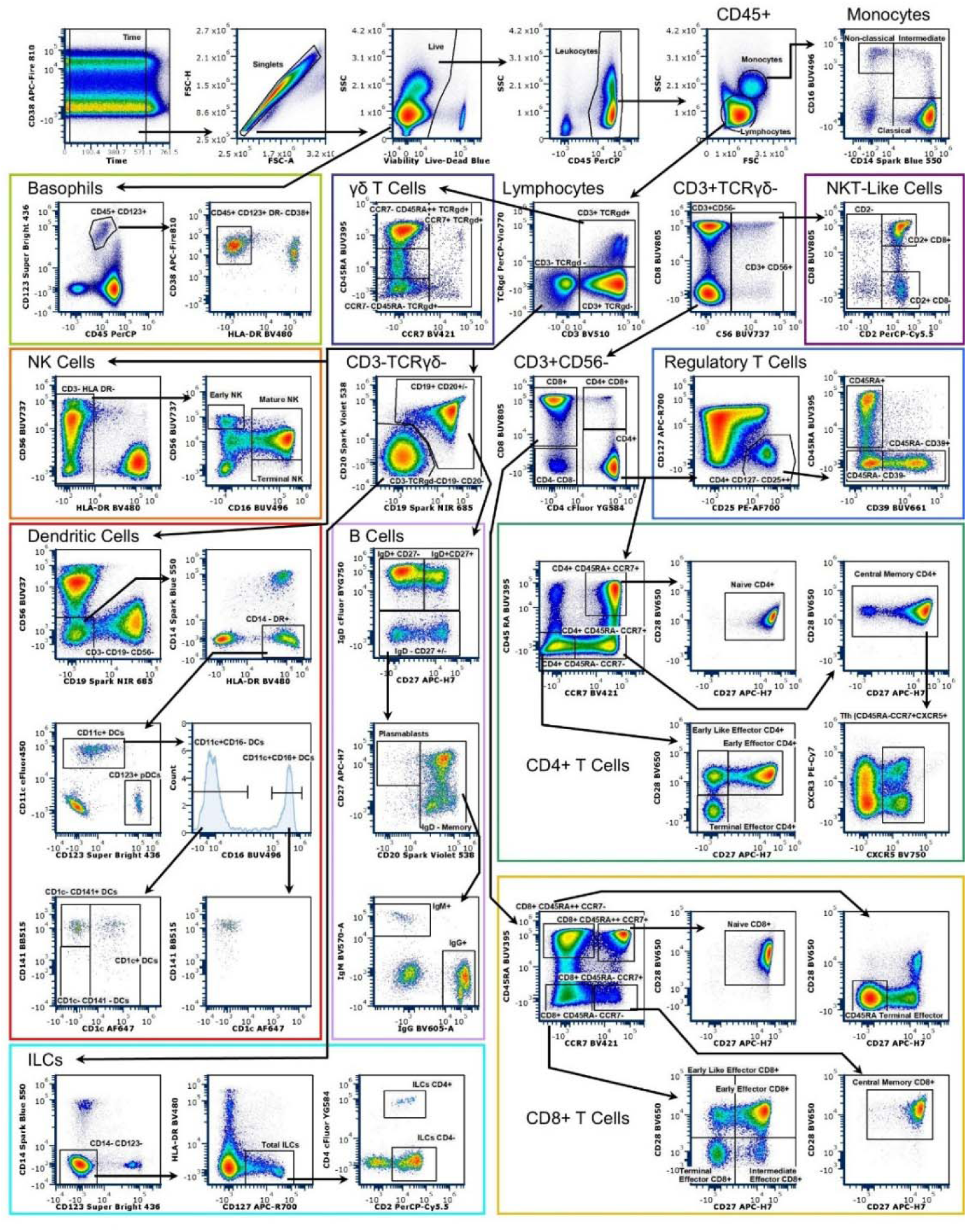
This 40-color panel characterizesCD4 T cells, CD8 T cells, regulatory T cells, γδ T cells, NKT-like cells, B cells, NK cells, monocytes, and dendritic cells. Below is the gating strategy used for this manuscript which follows the gating delineated by Park et. al ^10^.

Initial exploratory comparison of circulating immune populations across FeNO-defined phenotypes identified candidate immune subsets including classical monocytes, intermediate monocytes, and CCR7+ γδ T cells. To evaluate whether these candidate populations demonstrated relationships with FeNO as a continuous measure of airway inflammation, Spearman correlation analyses were performed. The distribution of these candidate populations was further characterized across healthy controls, FeNO-low asthma, and FeNO-high asthma.

In asthma, classical monocytes and non-classical monocytes exhibited a positive correlation with exhaled nitric oxide (classical monocytes: ρ = 0.52, p =0.029 and non-classical monocytes: ρ = 0.3, 0 =0.234) (Supplementary Figure 1). CCR7+ γδ T cells exhibited a negative correlation with exhaled nitric oxide (CCR7+ γδ T: ρ= -0.64, p= 0.004) (Supplementary Figure 1). To assess overall group differences within these subsets, Kruskal-Wallis test and subsequent Dunn’s post-hoc comparisons were applied across the immune subsets evaluated (Supplementary Table 2 and 3).

To visualize differences across FeNO-defined groups, candidate immune populations were displayed using boxplots, with statistical comparisons reported after BH adjustment. (Supplementary Table 4). FeNO-high asthmatics had higher classical (CD14++CD16−) and intermediate (CD14++CD16+) monocytes than healthy controls, (p = 0.025) (Figure 2). Nonclassical monocytes (CD14+CD16++) tended to be lowest in the low-FeNO group (Figure 2). In addition, FeNO-low asthma was distinguished by a unique subset of circulating T cells (with γδ receptors) that express CCR7 (C-C chemokine receptor 7) (Figure 2). These γδ T cells bridge innate and adaptive immunity and serve memory functions. Median proportions of CCR7+ γδ T cells were significantly higher in FeNO-low compared to FeNO-high asthma (Figure 2, all p < 0.01).

**Figure 2.**
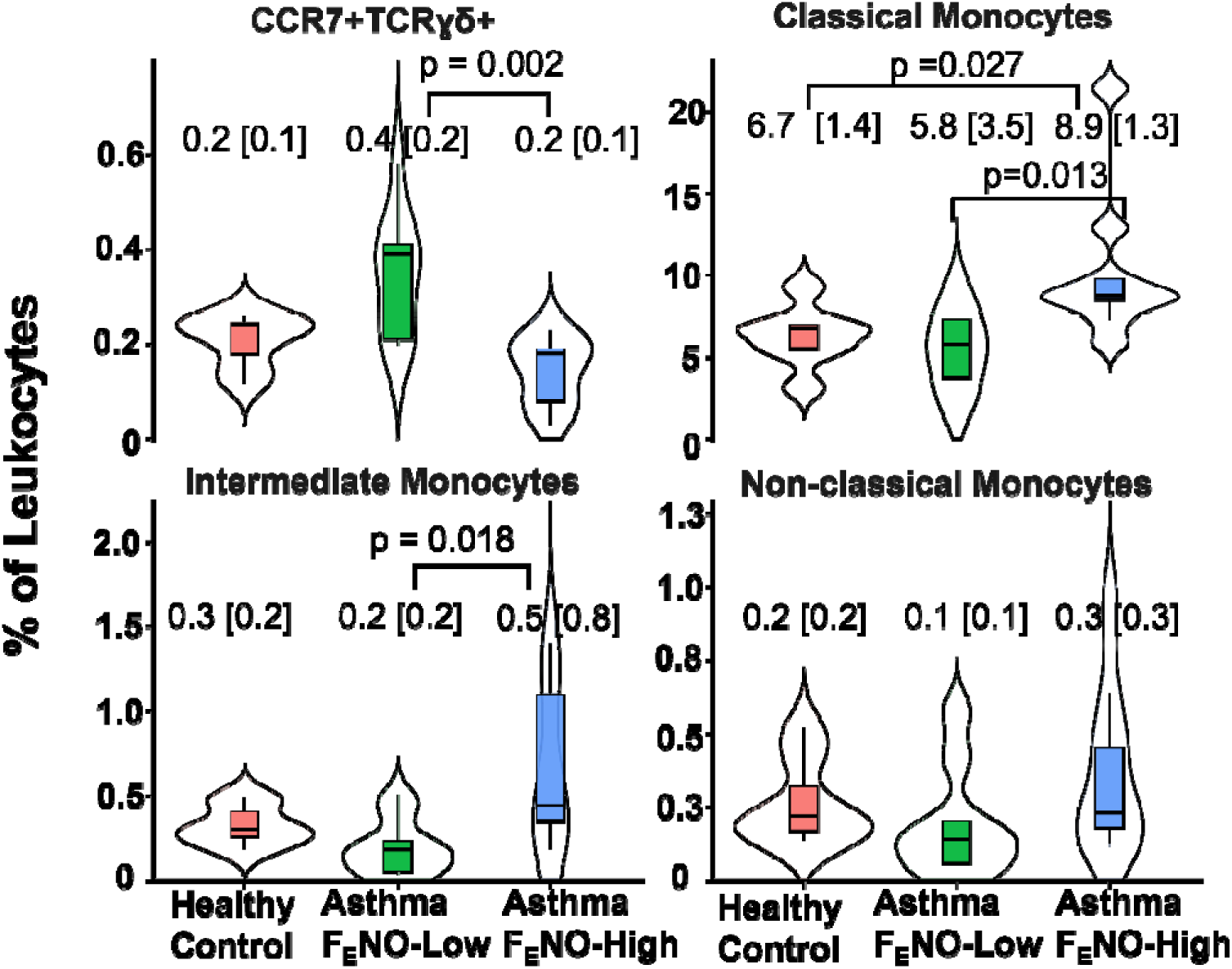
Participants with FeNO-high asthma exhibited higher proportions of circulating classical and intermediate monocytes, whereas FeNO-low asthma was associated with higher proportions of CCR7+ γδ T cells. The figure shows circulating leukocyte subset frequencies (expressed as a percentage of total leukocytes) among healthy controls, participants with FeNO-low asthma (<25 ppb), and participants with FeNO-high asthma (≥25 ppb). Horizontal lines indicate the median, and boxes represent the interquartile range (IQR). Overall group differences were assessed using the Kruskal–Wallis test. Pairwise comparisons were performed using Dunn’s test with Benjamini–Hochberg (BH) false discovery rate (FDR) correction for multiple comparisons.

Candidate FeNO-associated immune subsets identified through the exploratory immunophenotyping workflow were evaluated using Firth penalized logistic regression as a supportive analysis. Higher classical and intermediate monocyte frequencies were associated with increased odds of FeNO-high asthma, whereas CCR7+ γδ T-cell frequency was associated with decreased odds. Forest plot visualization of the model estimates demonstrated associations between FeNO status and several circulating immune cell populations (Figure 3). In the exploratory Firth logistic regression analyses, candidate immune subsets demonstrated that classical (CD14++CD16−) monocytes were associated with increased odds of high FeNO (OR 16.06, 95% CI 1.56–1304.69; p = 0.007), while intermediate (CD14++CD16+) monocytes also demonstrated a positive association with high FeNO (OR 4.19, 95% CI 1.22–132.30; p = 0.017). In contrast, CCR7+ γδ T cells were negatively associated with high FeNO (OR 0.06, 95% CI 0.00– 0.47; p = 0.001). Although the odds ratio for classical (CD14++CD16−) and intermediate monocytes (CD14++CD16+) demonstrated associations, the wide confidence intervals indicate substantial uncertainty in the estimate, likely due to the small sample size. o To evaluate the influence of individual observations in this limited sample, leave-one-out sensitivity analyses were performed. Sequential removal of each participant demonstrated preservation of the direction of association for classical monocytes, intermediate monocytes, and CCR7+ γδ T cells. This supports that the findings were not driven by a single influential participant. (Supplementary Table 5).

**Figure 3.**
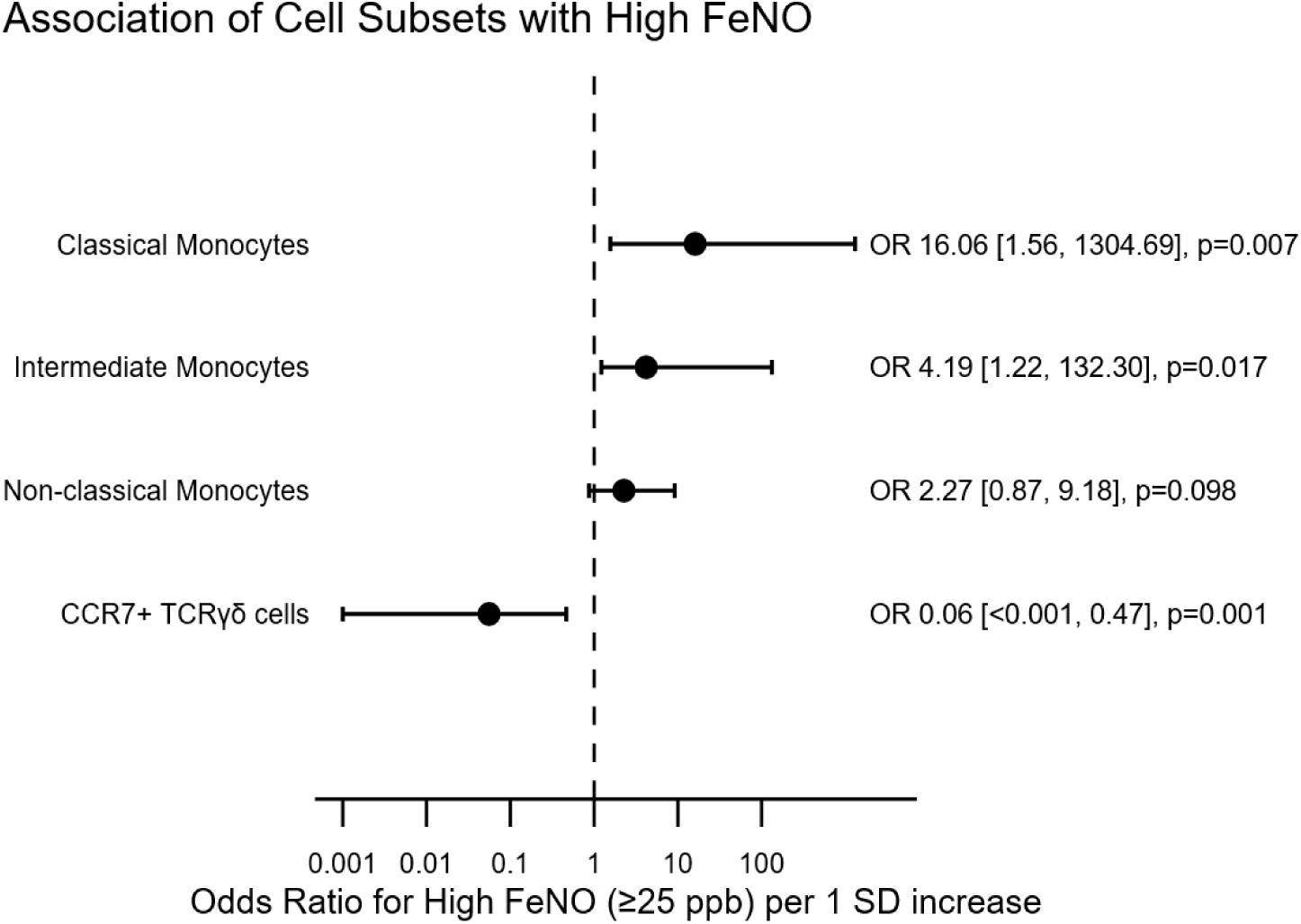
Forest plot of circulating immune cell subsets associated with FeNO-defined groups. Associations between immune cell subsets and elevated FeNO (≥ 25 ppb). Odds ratio (ORs) and 95% confidence intervals (CIs) represent the change in odds of high FeNO per 1 standard deviation increase in cell subset frequency, estimated using Firth logistic regression.

## Discussion

High-dimensional spectral flow cytometry enables detailed resolution of immune heterogeneity at the single-cell level, yet its application to asthma remains limited. To our knowledge, this is the first study employing spectral flow cytometric immunophenotyping to identify systemic immune signatures linked to FeNO-defined asthma phenotypes. Our results demonstrate distinct immune profiles in FeNO-high versus FeNO-low asthma, supporting the view the hypothesis these phenotypes are associated with distinct systemic inflammatory profiles. Specifically, FeNO-high asthmatics were associated with higher classical (CD14++CD16-) and intermediate monocytes (CD14++CD16+), while FeNO-low asthmatics were associated with higher CCR7+ γδ T cells and elevated TNF-α. In exploratory Firth logistic regression analyses, increased classical and intermediate monocyte frequencies were associated with greater odds of FeNO-high asthma, whereas increased CCR7+ γδ T-cell frequencies were associated with lower odds of FeNO-high asthma. The extent to which FeNO-defined asthma groups correspond to systemic immune phenotypes has not been fully understood, and differences in immunophenotype across immune cell subsets stratified by FeNO have not been previously reported ^3,4,21^.

Classical monocytes originate in the bone marrow and sequentially mature into intermediate monocytes and subsequently into non-classical monocytes in circulation. In healthy individuals, classical monocytes comprise 80-90% of circulating monocytes^22^. Classical and intermediate monocytes respond to and participate in type 2 inflammation. Classical monocytes are associated with more severe asthma compared to non-severe asthma ^23^.They have traditionally been associated with acute inflammatory recruitment, eosinophilic inflammation, airway infiltration, and cytokine production. As classical monocytes mature into intermediate monocytes (CD14++CD16+), they acquire enhanced antigen-presenting and pro-inflammatory capabilities, positioning them at the interface of the innate and adaptive immune response. Additionally, intermediate monocytes may contribute to angiogenic vasculogenesis through production of mediators involved in vasculogenesis because they produce mediators that drive the angiogenic switch in asthmatic airways^24,25,26^. Therefore, the observed circulating increase in classical and intermediate monocytes may reflect alterations in monocyte maturation or activation pathways associated with chronic inflammatory and vascular activation in asthma.

γδ T cells comprise approximately 5% of peripheral T cells and can exhibit diverse immunoregulatory and pro-inflammatory functions ^27^. We find that FeNO-low asthma, when compared to FeNO-high asthma, has a higher percentage of circulating CCR7+ γδ T cells, a distinct γδ T subset. FeNO-low asthmatics show the same trend as healthy controls, though this is not statistically significant. This increase suggests that CCR7+ γδ T cells may represent a candidate immune feature associated with FeNO-low asthma.

The specific role of CCR7+ γδ T cells in FeNO-low asthma remains largely unexplored in the literature. Interestingly, T-cell immunity pathways are enriched in type 2-low asthma through IL-17 associated inflammatory responses characteristic of T2-low disease. ^28,29^. However, γδ T cells have context-dependent functions and may contribute to either inflammatory or immunosuppressive immune responses.^29^ Experimental models have demonstrated that γδ T-cells can influence airway hyperresponsiveness, antiviral immunity, and cytokine production. Whether the increased circulating CCR7+ γδ T-cells observed in FeNO-low asthma contribute to these processes in human remains unknown ^30,28^.

The present study has several limitations, including the modest sample size. Given the small group sizes and the high-dimensional nature of the spectral flow cytometry data, the statistical analyses were designed to identify candidate immune populations for future validation rather than establish definitive biomarkers. Although several associations demonstrated relatively large estimated effect sizes, the corresponding confidence intervals were wide, reflecting the limited precision inherent to the sample size. Accordingly, these findings should be interpreted as hypothesis-generating and provide a framework for future investigation. In addition, the cross-sectional design precludes determination of causal relationships between circulating immune populations and airway inflammatory activity. Longitudinal studies will be important to determine whether immune signatures associated with FeNO-defined asthma phenotypes remain stable over time and predict clinical outcomes.

Participants with asthma had a higher BMI than healthy controls. Although this represents a limitation, obesity and metabolic dysfunction are well-established features of asthma and are commonly observed in real-world asthma cohorts. Consequently, the BMI distribution in our study is consistent with clinical practice and may enhance the generalizability of our findings. Another limitation is the potential influence of inhaled corticosteroids and biologic therapies on circulating immune populations. Therefore, these findings should be interpreted as reflecting the systemic immune signatures of treated asthma rather than untreated disease. Nevertheless, the persistence of distinct immune phenotypes despite contemporary treatment suggests that important systemic immunologic differences remain detectable in patients with asthma.

A strength of this study is the application of the OMIP-069 panel, as it provides a standardized 40-color spectral flow cytometry framework for deep immunophenotyping of peripheral blood immune populations for individuals with asthma. This approach enables simultaneous interrogation of diverse immune cell subsets and supports reproducible high-dimensional immune profiling across studies with a single acquisition.

The observed systemic immune cell subset signatures were associated with FeNO, suggesting a relationship between circulating immune phenotypes and airway inflammation. Whether these circulating immune populations directly contribute to airway inflammation or represent downstream consequences remains to be determined. These findings can be used for further mechanistic studies. The role of classical and intermediate monocytes should be further investigated in FeNO-high asthma. The role of γδ T cells also needs further elucidation. FeNO is used to guide asthma management, specifically biologic agents ^31^. A better understanding of the mechanisms underlying FeNO-low and FeNO-high asthma will help clinicians manage severe asthma more effectively.

In conclusion, our findings provide evidence that FeNO-defined asthma groups may be associated with distinct systemic immune signatures and warrant validation in larger studies. FeNO-high asthma was associated with elevated levels of classical and intermediate monocytes, consistent with enhanced systemic inflammation, whereas low-FeNO asthma shows increased CCR7⁺ γδ T cells. These findings suggest that airway inflammation, as reflected by FeNO, is accompanied by distinct systemic immune phenotypes and provide a foundation for future mechanistic investigations. FeNO-high asthma was associated with increased classical and intermediate monocytes, consistent with enhanced systemic inflammatory activity, whereas FeNO-low asthma was characterized by increased CCR7+ γδ T cells.

## Supporting information

Supplementary index

## Acknowledgments

**The authors would like to thank the clinical team, which includes Monica Labadia, Kevin Smith, and Michelle Koo,** for their assistance with recruitment. We also thank Joel Crespo (Cytek Biosciences) for his technical assistance with fine-tuning our Aurora spectral analyzer for OMIP-69. We thank the Cleveland Clinic Flow Cytometry Core Facility for excellent technical support with instrument QC and QA.

