## Supplementary index for "High-Dimensional Immunophenotyping Identifies Circulating Immune Signatures Associated with FeNO-Defined Asthma Phenotypes"

**Table 1. Reagents used for OMIP-069.** This 40-color fluorescent panel using full spectrum flow cytometry presents an in-depth characterization of lymphocytes, monocytes, and dendritic cells present in human peripheral blood. This panel covers major immune subsets of the human peripheral immune system. This table includes a final panel with cell marker with purpose, fluorochrome, and clone.

Abbreviations: APC = Allophycocyanin; PE = Phycoerythrin; PerCP = Peridinin-Chlorophyll-Protein.

| Marker | Fluorochrome | Clone | Purpose |
| --- | --- | --- | --- |
| Viability | Live-Dead Blue | NA | Live cells |
| CD4 | cFluor YG584 | SK3 | CD4 T cell, NKT-Like cell |
| CD16 | BUV 496 | 3G8 | Monocyte, NK cell, and dendritic cell differentiation |
| CD95 | PE-Cy5 | DX2 | T cell and B cell differentiation |
| CD8 | BUV 805 | SK1 | CD8 T, NK, and NKT-Like cells |
| CD19 | Spark NIR 685 | HIB19 | B cells |
| CD141 | BB515 | 1A4 | Dendritic cell differentiation |
| CD45RA | BUV 395 | SH9 | T cell and dendritic cell differentiation |
| HLA-DR | BV480* | L203 | T cell and monocyte activation, NK cell lineage discrimination, dendritic cell lineage marker |
| CD56 | BUV 737 | NCAM 16.2 | Pan NK cell, γδ T cell activation |
| CD159a | APC | REA110 | NK, NKT-Like, and γδ T cell activation/differentiation |
| CD159c | PE | REA205 | NK cell differentiation |
| CD45 | PerCP | HI30 | Leukocytes |
| CD14 | Spark Blue 550 | 63D3 | Monocyte Differentiation |
| IgM | BV570 | MHM-88 | B cell differentiation |
| CD337 | PE-Dazzle 594 | P30-15 | NK cell differentiation |
| CD38 | APC-Fire 810 | HIT2 | Monocyte, dendritic cell, T cell, and B cell activation/differentiation |
| CD20 | Spark Violet 538 | 2H7 | B cells |
| CD28 | BV 650 | CD28.2 | T cell and NK cell differentiation |
| CD3 | BV 510 | SK7 | Pan T cell, NKT-Like cells |
| CD24 | PE-Alexa Fluor 610 | SN3 | B cell differentiation |
| CD39 | BUV 661 | TU66 | B cell, T_regs_, and monocyte differentiation |
| CD2 | PerCP-Cy5.5 | TS1/8 | NK cell differentiation |
| CCR6 | BV711 | G034E3 | Chemokine receptor; T cell and B cell differentiation |
| CD314 | BUV 615 | 1D11 | NK cell differentiation |
| CD25 | PE-AlexaFluor 700 | CD25-3G10 | Regulatory T cells |
| CD27 | APC-H7 | M-T271 | T and B cell differentiation |
| CD127 | APC-R700 | HIL-7R-M21 | Cytokine receptor; T cell differentiation |
| IgD | cFluor BYG 750 | IgD26 | B cell differentiation |
| CD57 | cFluor B532* | HNK-1 | NK and CD8+ T cell immune senescence |
| CD1c | Alexa Fluor 647 | L161 | Dendritic cells, NKT-Like cells |
| CCR5 | BV 750 | RF8B2 | Chemokine receptor; monocyte, dendritic cell, T- cell, and B- cell differentiation |
| CD123 | Super Bright 436 | 6H6 | Plasmacytoid dendritic cells |
| IgG | BV 605 | G18-145 | B cell differentiation |
| TCR-ɣꟘ | PerCP-Vio 700 | REA591 | Pan γδ T cell |
| CXCR5 | BV 750 | RF8B2 | Chemokine receptor; T cell differentiation |
| PD-1 | BV 785 | BV785 | T cell inhibitory receptor |
| CXCR3 | PE-Cy7 | G025H7 | Chemokine receptor; Dendritic cell, T cell, and B cell differentiation |
| CCR7 | BV 421 | G043H7 | T cell differentiation |
| CD11c | eFluor450 | 3.9 | Dendritic cell differentiation |

**Table 2.** Deep immunophenotyping using OMIP-069 identified immune cell subsets that were subsequently analyzed according to FeNO-high and FeNO-low status. The numbers are proportions of cell populations compared to the total number of leukocytes acquired. Flow cytometry subset frequencies were analyzed using available case analysis. Lymphocytes and monocytes are included to provide context for downstream lymphocyte subset analyses. CD3+TCRgd- is a heterogeneous residual population. For each immune subset, participants with missing measurements were excluded from calculations of summary statistics and hypothesis testing for that specific subset. Continuous variables are presented as median (IQR), with corresponding p-values reported. Two asterisks signify unadjusted p value < 0.05. Abbreviations: BH = Benjamini-Hochberg; FeNO = exhaled nitric oxide.

| **Cellular Subset** | **FeNO-low** | **FeNO-high** | **p-value** | **BH-adjusted p-value** |
| --- | --- | --- | --- | --- |
| **n** | **10** | **8** |  |  |
| **Leukocytes** | | | | |
| **Lymphocytes** | 90.03 [88.35, 92.68] | 85.91 [82.20, 86.87] | 0.008 | 0.134 |
| **Monocytes** | 6.62 [4.71, 8.41] | 10.77 [9.5, 12.06] | 0.004 | 0.089 |
| **Basophils** | 0.20 [0.12, 0.38] | 0.41 [0.30, 0.56] | 0.130 | 0.462 |
| **Monocytes** | | | | |
| **Classical** | 5.79 [5.39, 7.32] | 8.80 [7.89, 12.16] | 0.004 | 0.089 |
| **Intermediate** | 0.18 [0.11, 0.38] | 0.92 [0.48, 1.33] | 0.010 | 0.134 |
| **Non-Classical** | 0.18 [0.10, 0.31] | 0.44 [0.36, 0.57] | 0.075 | 0.447 |
| **Tgd** | | | | |
| **CD3+ TCRgd+** | 1.80 [1.15, 3.52] | 1.61 [1.23, 2.29] | 0.424 | 0.743 |
| **Tgd-** | | | | |
| **CD3+ TCRgd-** | 70.13 [67.28, 74.66] | 59.61 [54.73, 66.58] | 0.026 | 0.249 |
| **CD3-** | | | | |
| **CD3- TCRgd -** | 17.46 [15.30, 18.36] | 17.93 [15.79, 24.36] | 0.374 | 0.743 |
| **Tgd (CD3+TCRgd+)** | | | | |
| **CCR7- CD45RA++ TCRgd+** | 0.28 [0.11, 0.48] | 0.50 [0.23, 1.22] | 0.594 | 0.847 |
| **CCR7+ TCRgd+** | 0.35 [0.21, 0.41] | 0.15 [0.08, 0.18] | 0.001 | 0.067 |
| **CCR7- CD45RA- TCRgd+** | 0.45 [0.14, 0.81] | 0.32 [0.11,0.51] | 0.398 | 0.743 |
| **T Cells CD3+TCRgd-** | | | | |
| **CD3+ CD56+** | 3.06 [1.64, 10.95] | 0.62 [0.45, 0.96] | 0.051 | 0.380 |
| **CD3+ CD56-** | 64.75 [57.10, 67.62] | 67.92 [53.90, 66.11] | 0.328 | 0.743 |
| **NKT (CD3+TCRgd-CD56+)** | | | | |
| **CD2-** | 0.03 [0.01, 0.06] | 0.03 [0.02, 0.05] | 0.821 | 0.952 |
| **CD2+ CD8+** | 1.76 [0.82, 7.74] | 0.45 [0.35, 0.78] | 0.91 | 0.447 |
| **CD2+ CD8-** | 0.24 [0.12.0.75] | 0.06 [0.04,0.10] | 0.013 | 0.145 |
| **T cells** | | | | |
| **CD8+** | 14.91 [13.42,17.32] | 13.05[8.78, 19.5] | 0.477 | 0.743 |
| **CD4+** | 44.66[38.94, 48.08] | 38.11 [37.03, 50.23] | 0.477 | 0.743 |
| **CD4+ CD8+** | 0.38 [0.27,0.48] | 0.28 [0.19,0.34] | 0.248 | 0.710 |
| **CD4- CD8-** | 1.14 [0.77, 1.74] | 0.88 [0.65, 1.05] | 1.31 | 0.468 |
| **Treg (CD3+CD4+TCRgd-CD56-)** | | | | |
| **CD4+ CD127- CD25++** | 2.50 [2.06, 3.34] | 2.98 [2.22, 3.13] | 0.965 | 0.995 |
| **CD45RA+** | 0.79 [0.51,1.17] | 0.72[0.56, 0.99] | 0.424 | 0.743 |
| **CD45RA- CD39+** | 0.94[0.62,0.99] | 1.08[0.77, 1.46] | 0.248 | 0.710 |
| **CD45RA- CD39-** | 0.50 [0.37, 0.90] | 0.60 [0.48, 0.68] | 0.929 | 0.988 |
| **CD4+ T Cells (CD3+ CD4+ TCRgd- CD56-)** | | | | |
| **CD4+** | 44.66 [38.94, 48.08] | 38.11 [37.03, 50.23] | 0.477 | 0.743 |
| **CD4+ CD45RA+ CCR7+** | 21.68 [15.48, 29.77] | 19.23[ 17.04, 24.68] | 0.929 | 0.988 |
| **Naive CD4+** | 21.63 [15.43, 29.74] | 19.22[16.99, 24.65] | 0.965 | 0.995 |
| **CD4+ CD45RA- CCR7+** | 11.81[9.09, 13.63] | 10.96 [10.02, 16.82] | 0.722 | 0.896 |
| **Central Memory CD4+** | 11.80 [9.07, 13.62] | 10.92 [10.01, 16.81] | 0.722 | 0.896 |
| **CD4+ CD45RA- CCR7-** | 7.04 [5.29, 11.38] | 7.98 [2.74, .51] | 0.722 | 0.896 |
| **Early Like Effector CD4+** | 1.47 [1.02, 2.27] | 1.31 [1.04, 1.85] | 0.859 | 0.959 |
| **Early Effector CD4+** | 4.6 [3.42, 9.00] | 4.44 [3.07, 6.066] | 0.534 | 0.795 |
| **Terminal Effector CD4+** | 0.03 [0.02, 0.23] | 0.05 [0.01,0.34] | 1.00 | >0.999 |
| **CD8+ T Cells (CD3+ CD8+ TCRgd- CD56-)** | | | | |
| **CD8+** | 14.91 [13.42, 17.32] | 13.05 [8.78, 19.95] | 0.477 | 0.743 |
| **CD8+ CD45RA++ CCR7+** | 6.20 [3.46, 7.35] | 5.18 [1.23, 7.56] | 0.424 | 0.743 |
| **Naive CD8+** | 6.17 [3.43, 7.32] | 5.16 [1.23, 7.56] | 0.424 | 0.743 |
| **CD8+ CD45RA- CCR7+** | 0.77 [0.61, 1.10] | 0.87 [0.69, 1.00] | 0.722 | 0.896 |
| **Central Memory CD8+** | 0.76 [0.58, 1.09] | 0.86 [0.67,0.98] | 0.657 | 0.896 |
| **CD8+ CD45RA++ CCR7-** | 3.80 [1.06,6.43] | 1.67 [1.12, 4.43] | 0.790 | 0.945 |
| **CD45RA+ Terminal Effector CD8+** | 0.54 [0.30, 4.03] | 0.91 [0.36, 2.56] | 0.929 | 0.988 |
| **CD8+ CD45RA- CCR7-** | 2.26 [0.95,3.58] | 2.97 [1.58, 3.72] | 0.477 | 0.743 |
| **CD28+CD27-** | 0.28 [0.07, 0.40] | 0.29 [0.18, 0.44] | 0.328 | 0.743 |
| **CD28+CD27+** | 1.54 [0.60, 2.22] | 1.44 [0.99, 2.53] | 0.534 | 0.795 |
| **CD28-CD27+** | 0.17 [0.11, 0.30] | 0.30 [0.22,0.39] | 0.230 | 0.710 |
| **CD28-CD27-** | 0.11 [0.06,0.36] | 0.32 [0.14, 0.73] | 0.214 | 0.710 |
| **NK Cells** | | | | |
| **Early NK** | 0.36 [0.21, 0.43] | 0.36 [0.33, 0.40] | 1.000 | >0.999 |
| **Mature NK** | 4.72 [3.17, 6.09] | 7.38 [4.20, 8.48] | 0.051 | 0.380 |
| **Terminal NK** | 0.24[0.13,0.35] | 0.20 [0.17, 0.28] | 0.859 | 0.959 |
| **Dendritic Cells** | | | | |
| **CD11c+ DCs** | 0.31 [0.18,0.58] | 0.42 [0.26,0.79] | 0.657 | 0.896 |
| **CD11c+ CD16+** | 0.08 [0.03, 0.37] | 0.20 [0.07, 0.34] | 0.449 | 0.743 |
| **CD11c+ CD16-** | 0.17 [0.11, 0.28] | 0.20 [0.17, 0.31] | 0.261 | 0.710 |
| **CD1c+ DCs** | 0.08 [0.04, 0.12] | 0.10 [0.03, 0.11] | 0.824 | 0.952 |
| **B Cells (CD3- CD19+ CD20+/-)** | | | | |
| **CD1c- CD141+ DCs** | 0.17 [0.11,0.28] | 0.20 [0.17, 0.31] | 0.265 | 0.710 |
| **CD123+ pDCs** | 0.04 [0.03,0.50] | 0.08 [0.04, 0.15] | 0.089 | 0.447 |
| **CD1c- CD141 - DCs** | 0.00 [0.00, 0.01] | 0.01 [0.01, 0.01] | 0.392 | 0.743 |
| **CD19+ CD20+/-** | 7.94 [5.62, 9.22] | 6.18 [4.45, 8.59] | 0.424 | 0.743 |
| **IgD+ CD27-** | 6.24 [4.26, 7.16] | 4.64 [2.96, 7.55] | 0.722 | 0.896 |
| **IgD+CD27+** | 0.46 [0.32, 0.59] | 0.35 [0.22, 0.48] | 0.328 | 0.743 |
| **IgD - CD27 +/-** | 1.04 [0.94, 1.57] | 0.79 [0.65, 1.04] | 0.100 | 0.447 |
| **IgD - Memory** | 0.95 [0.84, 1.37] | 0.74 [0.60, 0.90] | 0.091 | 0.447 |
| **IgM+** | 0.04 [0.03, 0.08] | 0.04 [0.02, 0.06] | 0.590 | 0.847 |
| **IgG+** | 0.61 [0.46, 0.70] | 0.41 [0.36, 0.45] | 0.100 | 0.447 |
| **Plasmablasts** | 0.04 [0.03, 0.07] | 0.03 [0.02, 0.06] | 0.323 | 0.743 |
| **ILCs** | | | | |
| **Total ILCs** | 0.56 [0.38, 0.66] | 0.41 [0.36, 0.44] | 0.131 | 0.462 |
| **ILCs CD4+** | 0.04 [0.02, 0.06] | 0.03 [0.03, 0.04] | 0.782 | 0.945 |
| **ILCs CD4-** | 0.28 [0.21, 0.38] | 0.20 [0.12, 0.24] | 0.109 | 0.456 |

Figure 1. Relationship between exhaled nitric oxide and monocyte and γδ T-cell subsets in asthma. Scatterplots show associations between FeNO and the frequencies of classical, intermediate, and non-classical monocytes, as well as CCR7+ γδ T cells. These populations were selected for visualization because exploratory analyses identified coordinated associations within monocyte and γδ T-cell compartments. Spearman correlation coefficients (ρ) and p-values are displayed in each panel.

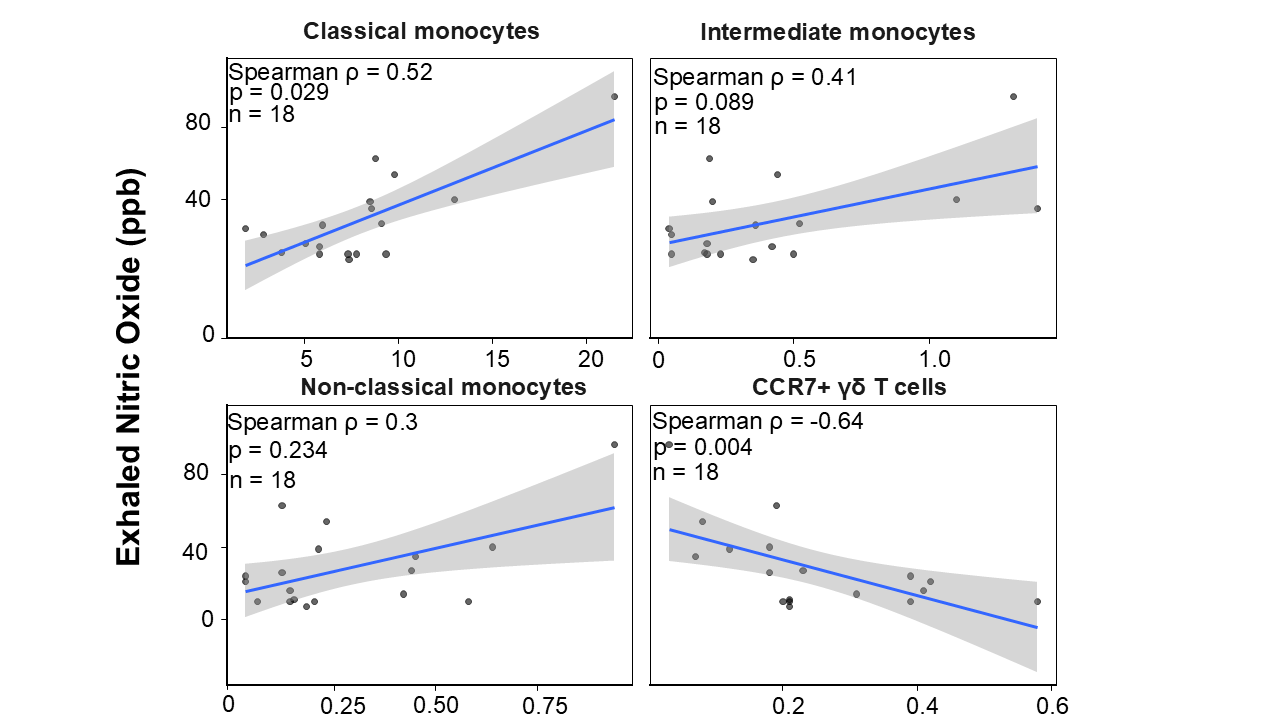

**Table 3.** Overall differences in immune cell subset frequencies across healthy controls, FeNO-low asthma, and FeNO-high asthma groups were assessed using the Kruskal–Wallis test with Benjamini–Hochberg false discovery rate correction. Significant overall differences were observed for CCR7+ γδ T cells (adjusted *P* = 0.0098), classical monocytes (adjusted *P* = 0.0196), and intermediate monocytes (adjusted *P* = 0.0285), whereas nonclassical monocytes were not significantly different among groups (adjusted *P* = 0.132).

| Subset | P-value | BH-adjusted p-value |
| --- | --- | --- |
| **CCR7+ γδ T cells** | **0.0025** | **0.0098** |
| **Classical monocytes** | **0.0098** | **0.0196** |
| **Intermediate monocytes** | **0.0214** | **0.0285** |
| Nonclassical monocytes | 0.132 | 0.132 |

**Table 4.** Participants with FeNO-high asthma exhibited higher proportions of circulating classical and intermediate monocytes, whereas FeNO-low asthma was associated with higher proportions of CCR7+ γδ T cells. The figure shows circulating leukocyte subset frequencies (expressed as a percentage of total leukocytes) among healthy controls, participants with FeNO-low asthma (<25 ppb), and participants with FeNO-high asthma (≥25 ppb). After overall group differences were assessed using the Kruskal–Wallis test, pairwise comparisons were performed using Dunn's test with Benjamini–Hochberg (BH) false discovery rate (FDR) correction for multiple comparisons.

| Immune Subset | Comparison | N | N | p-value | BH-adjusted p-value |
| --- | --- | --- | --- | --- | --- |
| CCR7+γδ+ | Healthy vs FeNO-low asthma | 7 | 10 | 0.169 | 0.1700 |
| CCR7+γδ+ | Healthy vs FeNO-high asthma | 7 | 8 | 0.062 | 0.0936 |
| **CCR7+γδ+** | **FeNO-low versus FeNO-high asthma** | **10** | **8** | **0.0005** | **0.0016** |
| Classical Monocytes | Healthy versus FeNO-low asthma | 7 | 10 | 0.7933 | 0.7933 |
| **Classical monocytes** | **Healthy versus FeNO-high asthma** | **7** | **8** | **0.0181** | **0.0271** |
| **Classical Monocytes** | **FeNO-low versus FeNO- high asthma** | **10** | **8** | **0.0044** | **0.0131** |
| Intermediate monocytes | Healthy versus FeNO-low asthma | 7 | 10 | 0.1463 | 0.2195 |
| Intermediate Monocytes | Healthy versus FeNO-high asthma | 7 | 8 | 0.2533 | 0.2533 |
| **Intermediate Monocytes** | **Asthma Feno-low versus FeNO-high asthma** | **10** | **8** | **0.0059** | **0.0176** |
| Non-classical monocytes | Healthy versus FeNO-low asthma | 7 | 10 | 0.1862 | 0.2794 |
| Non-classical monocytes | Healthy versus FeNO-high asthma | 7 | 8 | 0.6054 | 0.6054 |
| Non-classical monocytes | FeNO-low versus FeNO-high asthm | 10 | 8 | 0.0528 | 0.1583 |

**Table 5. Leave-one-out sensitivity analysis of Firth logistic regression evaluating the association between CCR7+γδ T cells and FeNO-defined asthma phenotype.** Exploratory modeling demonstrated opposing immune signatures associated with FeNO-defined phenotypes, with FeNO-high asthma characterized by enrichment of circulating classical/intermediate monocytes and relative depletion of CCR7+ γδ T cells.

| **Candidate Immune Subset** | **Models refit** | **OR Direction maintained** | **Range of leave-one-out ORs** |
| --- | --- | --- | --- |
| Classical monocytes | 18 | Increased odds of FeNO-high in all models | 11.5–118.0 |
| Intermediate monocytes | 18 | Increased odds of FeNO-high in all models | 3.4–6.1 |
| CCR7+ γδ T cells | 18 | Decreased odds of FeNO-high in all models | 0.016–0.080 |
